# 3-epicaryoptin induces G2/M phase cell cycle arrest and apoptosis in human breast cancer cells by disrupting the microtubule network, an *in vitro* and *in silico* investigation

**DOI:** 10.1101/2024.01.04.574171

**Authors:** Manabendu Barman, Sujit Roy, Nanda Singh, Debanjan Sarkar, Niharendu Barman, Amit Pal, Sankar Bhattacharyya, Sanjib Ray

## Abstract

Breast cancer (BC) is a prevalent form of cancer observed in women across the globe, constituting over a quarter of all female BC cases. The treatment of BC continues to require significant efficacy, aiming to achieve high success rates while minimizing adverse effects on the body as a whole. In the current study, 3-epicaryoptin was tested for the molecular mechanism of its anti-cancer activity in the human breast cancer cell line, MCF-7. We investigated cell viability by MTT assay, cell cycle kinetics and apoptosis, immunofluorescence straining, molecular modelling, and ADMET profiling. MTT assay results showed that 3-epicaryoptin was found cytotoxic against MCF-7 cells with an IC_50_ value of 344.64 µg mL^-1^ for 48 h. Flow cytometric analysis exhibited that 3-epicaryoptin halted the MCF-7 cells in the G2/M phase and subsequently induced apoptosis in a time-dependent manner. Our immunofluorescence studies indicated that 3-epicaryoptin inhibited microtubule polymerization in MCF-7 cells. Furthermore, molecular docking followed by molecular dynamics (MD) simulation studies demonstrated the ability of 3-epicaryoptin to interact with the tubulin protein at the colchicine binding pockets. Overall, our results suggest that 3-epicaryoptin can inhibit the proliferation of human breast cancer cells by depolymerizing of cellular microtubule networks, which causes cell cycle arrest and promotes apoptotic cell death. Therefore, it has been indicated that the natural product 3-epicaryoptin exhibited considerable promise as a potent therapeutic agent capable of inducing apoptosis in breast cancer cells.

## Introduction

Breast cancer (BC) is a significant and widespread health issue, representing 14.7% of all cancer-related deaths in women [1]. The risk of BC substantially rises with age [2]. BCs account for over 40% of female malignancies diagnosed by age 40 and 20% before age 30 [3]. Various treatment options, including surgery, chemotherapy, and radiotherapy, are available for BC. However, each treatment method carries risks and adverse effects. Furthermore, several clinical drugs used in BC treatment exhibit notable side effects, which can undermine patient confidence in the therapeutic process. Thus, there is an urgent need to develop new and more efficient anticancer agents characterized by low toxicity to address BC.

Microtubules are the cytoskeletal protein filaments made up of α-and β-tubulin heterodimers [4]. It plays a very crucial role in various aspects of cellular processes, like in mitosis and cell division, for the maintenance of cell shape, vesicle transport, and cell motility [5]. Therefore, rendering these microtubule networks has become a promising target for the treatment of cancer [6]. In the last few decades, intense research has yielded a large number of microtubule targeting compounds that act as potent anticancer agents [7–10]. These compounds are mainly divided into two groups: the microtubule stabilizing agents (MSAs) and the microtubule destabilizing agents (MDAs). The terms microtubule ‘stabilizers’ or ‘destabilizes’ come from either the targeting compound increasing or decreasing the polymerization of tubulin [11–14]. Established MSA drugs include paclitaxel (the first identified agent in this class), docetaxel (Taxotere), epothilones, and discodermolide. Whereas the MDAs drugs include the compounds vinca alkaloids (vincristine, vinblastine, vinorelbine, vinflunine, and vindesine), podophyllotoxin, estramustine, colchicine, and combretastatins [15–18]. These antimitotic agents interfere with tubulin protein by binding to the taxol, vinblastine, or colchicine binding sites [5]. However, despite their therapeutic potential, microtubule inhibitors showed limitations such as the development of drug resistance due to frequently used, side effects, poor oral bioavailability, and low aqueous solubility [19, 20]. Therefore, the finding of new microtubule inhibitors with pharmacological efficacy and acceptable toxicological properties is required for the preclinical and clinical development.

3-epicaryoptin is a natural diterpenoid first isolated and identified by Hosozawa et al. (1974a) from the leaves of *Clerodendrum calamitosum* Maxim (Verbenaceae) [21]. Biological activity studies reported its potent insect antifeedant activity against various pests, including the tobacco cut worm and the potato beetle. Additionally, 3-epicaryoptin has shown inhibitory effects on the growth and mortality of European corn borer larvae and larvae of *Musca domestica* and *Culex quinquefasciatus* [22, 23, 21, 24].

In our previous comparative cytotoxicity analysis on a plant test system (*Allium cepa* root apical meristem cells) was shown that the compound 3-epicaryoptin isolated from the leaves of *C. inerme* has colchicine-like effects, including root tip swelling, metaphase arresting, micronuclei, and polyploidy inducing effects [25–29]. However, whether 3-epicaryoptin regulates the cell cycle in human breast carcinoma remains unclear. So, therefore, in the present investigation, we evaluate the *in vitro* cytotoxic effects of compound 3-epicaryoptin on MCF-7 cells and investigate the mechanistic actions leading to cancer cell death. Furthermore, to identify the possible molecular interaction of 3-epicaryoptin with the tubulin, we employed computational approaches involving molecular docking (MD), molecular dynamics simulation (MDS), and binding energy calculation studies.

## Materials methods

### Chemicals

Triton X-100, MTT reagent, and paraformaldehyde were obtained from Himedia, India. DMEM (Dulbecco’s Modified Eagle’s Medium), FBS (Fetal Bovine Serum), Penicillin-streptomycin, and 0.25% Trypsin-EDTA (1X) were obtained from Invitrogen-Gibco. RNase A, Propidium Iodide, DAPI/Antifade solution, and Annexin V-FITC apoptosis detection kit were purchased from Sigma-Aldrich, USA. Anti-tubulin antibody and Goat anti-mouse IgG-FITC used were obtained from Santa Cruz, the USA. 3-epicaryoptin was isolated from leaves of *Clerodendrum inerme* [26].

### Cell culture

Human breast cancer cell line, MCF-7, was purchased from the National Centre for Cell Science (Pune, India) and HiFi™ human peripheral blood mononuclear cells (H-PBMC) was purchased from HiMedia Laboratories Pvt. Ltd. Cells were cultured in Dulbecco’s MEM medium supplemented with 10% FBS, 100 µg mL^-1^ penicillin, 100 µg mL^-1^ streptomycin, and 2 mM L-Glutamine (Invitrogen-Gibco). Cells were incubated in humidified atmosphere with 5% CO_2_ at 37L.

### Study of cytotoxicity by MTT assay

The cytotoxic effect of 3-epicaryoptin was monitored by routine MTT colorimetric assay. Briefly, the MCF-7 cells and human PBMCs were seeded into a 24 well plate at a density of 5 × 10^3^ cells/well and incubated for 24 h at 37°C CO_2_ incubator in DMEM culture medium supplemented with 10% FBS, 100 µg mL^-1^ penicillin, 100 µg mL^-1^ streptomycin, and 2 mM L-Glutamine, respectively. Subsequently, cell line cultures were exposed to different concentrations (12.5–400 μg mL^-1^) of 3-epicaryoptin for a period of 24 and 48 h. After the treatment, MTT reagent (5 mg mL^-1^) was added to each well and then incubated at 37°C for 3 h. Next, the culture medium was discarded from each well and added DMSO into the well to dissolve the formed formazan crystals within metabolically live cells. Absorbance was measured at 570 nm on a microplate reader (Multiscan EX, Thermo scientific, USA) to determine the percentage of viable cells [30, 31].

### Cell cycle assay

The cell-cycle distribution was performed by flow cytometric DNA analysis. MCF-7 cells were cultured in 25 cm^2^ culture flasks at a density of 1 × 10^6^ cells/flask in DMEM medium supplemented with 10% FBS and antibiotic solution, kept at 37°C in a humidified atmosphere and 5% CO_2_ in the air. Upon 60-70 % of confluence, the cells were treated with 100 and 200 µg mL^-1^ concentrations of 3-epicaryoptin and incubated for 20 h. After treatment, the cells were harvested and washed with ice-cold PBS, fixed with chilled 70% ethanol (30 min at 4°C). The cells solution were then centrifuged to remove the ethanol, washed with cold PBS and added RNase (50 µg mL^-1^), incubated at 37 °C for 1 h, then stained with 50 µg mL^-1^ of propidium iodide (PI) solution for another 15-20 min by incubating at room temperature. Flow cytometric analysis of cell cycle was conducted using Beckman Coulter flow cytometer and the data were analyzed using CytExpert software, version 2.3 (Beckman, USA).

### Annexin V/PI apoptosis study

The Annexin V-FITC/PI staining assay was performed to determine the induction of apoptotic cell death. Seeded MCF-7 cells in 6-well culture plates were treated with 100 and 200 µg mL^-1^ concentrations of compound 3-epicaryoptin or vehicle (DMSO). After incubation for 24 and 48 h, treated and untreated cells were harvested, washed with ice-PBS, and incubated with Annexin V-FITC and PI. The samples were analyzed using a BD FACS Calibur flow cytometer (BD Biosciences, San Jose, CA, USA).

### Immunofluorescence staining

Immunofluorescence staining method was performed for the visualization of the effect of 3-epicaryoptin on cellular microtubules. MCF-7 cells (at a density of 1 × 10^5^ cells/mL) were grown on lysine coated glass coverslip in a 6 well cell culture plate, and then, upon 60-70 % of confluence, the cells were incubated with or without 100 and 200 µg mL^-1^ concentration of 3-epicaryoptin for 20 h. Afterward, the cells were subjected to fixation in paraformaldehyde (3.7 %) and then washed in PBS, permeabilization in 0.1% Trition X (10 min), and blocking of nonspecific binding with 1 % BSA for 2 h and then washed in PBS. The anti-β-tubulin primary antibody was used to probe the cellular microtubule and was incubated overnight and then washed with PBS, which was followed by incubation with FITC-conjugated goat anti-mouse IgG secondary antibody. DAPI was used to stain the nucleus. The images were captured using a Zeiss LSM 710, GmbH laser scanning confocal microscope (Carl Zeiss, Germany).

### Molecular modeling study

To identify the binding affinity of 3-epicaryoptin with tubulin, 3-epicaryoptin was docked onto the crystal structure of tubulin by using the molecular docking software AutoDock Vina v1.2.5 [32, 33]. The 3D crystal structure of αβ tubulin (Protein Data Bank ID: 1SA0.pdb) with a resolution of 3.58 Å was used as a receptor to perform the molecular docking study [13]. We chose only chain A and B of tubulin as a template and removed all the coordinates (ligand, GTP, GDP, and Mg^+2^) of chain A and B, other chains B & C together with the stathmin like domain from the crystal structure of 1SA0 using UCSF chimera version 1.14 [34, 35]. The 3D structure of 3-epicaryoptin was downloaded from the ZINC database and was ready to use [36]. In order to evaluate the quality of the docking protocol, DAMA-colchicine was extracted from the crystal structure and redocked into the binding site. Then the RMSD (Root Mean Square Deviation) value between re-docked ligand and co-crystallized conformation was calculated. After the parameters were verified by docking of known ligands (DAMA-colchicine), a similar method was carried out for the docking of 3-epicaryoptin on the tubulin dimer [37]. Initially, a blind docking of 3-epicaryoptin on tubulin dimer was performed. The input files such as protein and ligand pdbqt file as well as grid box coordinates were generated in AutoDock Tools 1.5.6 package [38]. The grid size was set to 102×86×62 xyz points with grid spacing of 1 Å. The grid center was set at dimensions (x, y, z); 120.224, 92.436, 10.560. The docking study were performed considering all rotatable angles of 3-epicaryoptin as flexible and tubulin as rigid. All parameters during docking were kept as default, with the exception of energy_range was set to 4 kcal, num_modes 20, and the value of exhaustiveness was set to 20. Ten independent docking runs were performed, and in each individual run, the best-ranked conformation according to the Vina docking score was found to be at the interface of the tubulin dimer. So, site-specific docking was carried out at the dimer interface.

For site-specific docking, we used a grid box that covered the interface of the αβ tubulin heterodimer, with dimensions of size_x: 40, size_y: 40, and size_z: 40, with a grid spacing of 1 Å. The grid was centered at X: 120.224, Y: 92.436, and Z: 10.560. During Vina docking, the tubulin heterodimer was kept as a rigid and 3-epicaryoptin as a flexible molecule. Other parameters, such as the value of exhaustiveness, were set to 200, energy_range = 4, and num_modes = 200. The best-ranked conformation as judged by the Vina docking score was selected and visually inspected using AutoDock 4.2 tools, UCSF Chimera version 1.14, and LigPlot^+^ version 2.2 program [32, 39, 38, 34, 33].

### Molecular dynamics (MD) simulation

The docked conformation of the complex with lowest binding energy was taken for Molecular Dynamics (MD) simulation to assess its structural stability and conformational flexibility. The MD simulation was performed using the GROMACS code (2021.4 version). The coordinates and topology files of the protein was prepared using the pdb2gmx module in GROMACS. The CHARMM36 force field [40–42] was adopted for the protein topology. The ligand topology file was generated using CHARMM General Force Field (CGenFF) [43, 44] server. The system was accommodated within a cubic box, maintaining a minimum distance of 1.0 nm from the edges, and solvated using the TIP3P water model. To neutralize the ligand-protein complex system, appropriate amount of Na^+^ and Cl^-^ were added as counter ions. Energy minimization of the molecular system was performed using the steepest descent method followed by the conjugate gradient method until the maximum force become less than 500 kJ mol^-1^ nm^-1^ [45]. After that, the system was equilibrated for 200 ps using NVT and the NPT ensemble protocol with 100000 steps. Simulations were conducted under stable conditions at a constant temperature of 300 K using the velocity-rescaling algorithm and constant pressure of 1 bar employing the Berendsen barostat algorithm [46]. The subsequent production run was carried out for 25 ns with trajectories generated for 2 fs time step [47]. Finally, the generated trajectories are used to obtain the root mean square deviation (RMSD), root mean square fluctuation (RMSF), solvent accessible surface area (SASA), and radius of gyration (rGy), employing Gromacs suite tools.

### MMPBSA calculations

The free energies (Δ*G*) was calculated using Molecular Mechanics/Poisson-Boltzmann Surface Area (MMPBSA) method [48], implemented in gmx_MMPBSA v1.6.2 code [49]. In this method, the binding free energy (Δ*G*_bind_) of the protein with ligand in solvent was expressed as

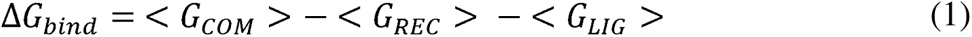

where, *G_COM_, G_REC_, G_LIG_* are the free energies of the complex, receptor and ligand, respectively which are given by

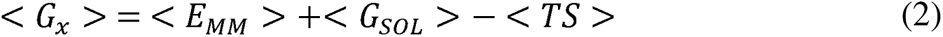

Here, *E_MM_* is the gas phase molecular mechanics energy, *G_SOL_* is the solvation energy term, *T* is the temperature and *S* is the entropy of the system.

Therefore, Δ*G_bind_* can be represented as,

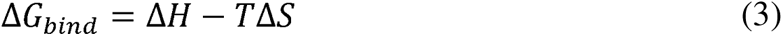

Where, Δ*H* == Δ*E_MM_* + Δ*G_SOLV_* is the enthalpy of binding. Δ*H* can further be decomposed as

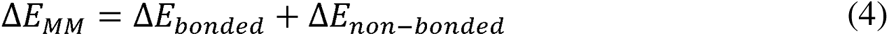

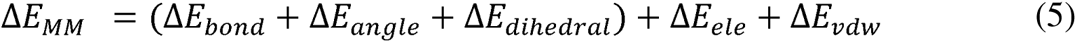

and

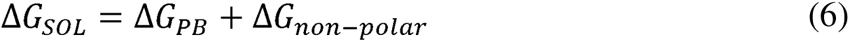

Here, Δ*E_bonded_* corresponds to the sum of bond, angle and dihedral contributions and Δ*E_non-bonded_* corresponds to the sum of electrostatic and van der Waals contribution to the Δ*E_MM_*. The polar contribution to solvation energy was calculated using Poisson-Boltzmann solvation model whereas the non-polar was calculated from the SASA.

In the present study, 300 frames are extracted from the last 15 ns MD trajectory was used to calculate the binding free energy. The entropy effect was neglected in the calculation.

### ADMET Prediction

We performed *in silico* ADME analysis to examine the physicochemical characteristics of the compound 3-epicaryoptin, including factors such as solubility, lipophilicity, oral acceptability, drug-likeness, and pharmacokinetics. The analysis was carried out using the online platform http://www.swissadme.ch [50]. However, the toxicity of these compound was not assessed through SwissADME; therefore, we utilized the pkCSM pharmacokinetics server to predict their toxicity properties based on their SMILE (simplified molecular input line entry specification) profile [51].

### Statistical analysis

Statistical analyses were performed using the OriginPro 8 software package. The level of significance at *p*≤ 0.05 or 0.01 or 0.001, between the control and treated values for the cell frequencies at different phases, and cellular viability were analyzed by Student’s *t*-test.

## Result

### MTT assay

The cellular toxicity of 3-epicaryoptin was assessed by performing the MTT assay on both the human cancer cell line MCF-7 and normal **PBMCs (Figure 1)**. The study results showed that the treatment of 3-epicaryoptin in MCF-7 cells causes a concentration-dependent decrease in the viable cell percentage after 24 and 48 h, respectively. Significantly (*p*< 0.001) highest cytotoxicity against MCF-7 cells was found at the highest treatment concentration (400 µg mL^-1^) of 3-epicaryoptin after 48 h, resulting in a viable cell percentage of 48.36±1.63 %. A significant (*p*< 0.001) reduction in the viable cell percentage was also observed after 24 h. The concentration of 3-epicaryoptin that reduced the survival of cells by 50% (IC_50_ values) was found to be 344.64 µg^-1^ mL after 48 h (**Figure 1A**). Conversely, treatment with 3-epicaryoptin on normal PBMCs did not exhibit significant cytotoxicity after 24 h (**Figure 1B**). These results provide additional proof of 3-epicaryoptin’s potential to preferentially kill cancerous MCF-7 cells compared to normal PBMCs.

**Fig. 1.**
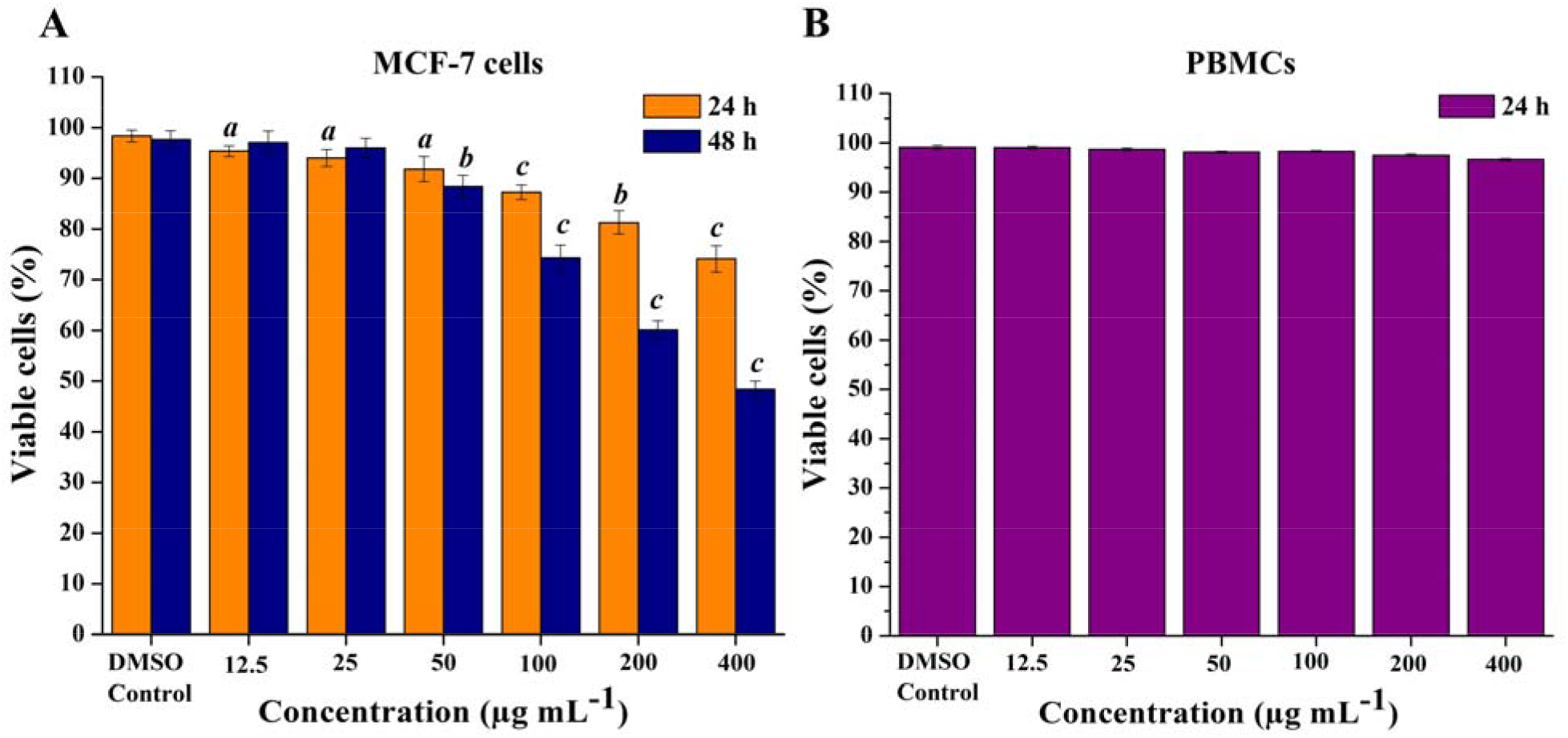
Cytotoxicity study of 3-epicaryoptin on MCF-7 cells and human peripheral blood mononuclear cells (PBMCs) by the MTT assay. (A) Shows the effect of 3-epicaryoptin (12.5-400 µg mL^-1^) on the percentage of viable cells evaluated by MTT assay in MCF-7 cells after 24 and 48 h treatment. (B) Shows the effect of 3-epicaryoptin (12.5-400 µg mL^-1^) on the percentage of viable cells evaluated by MTT assay in PBMCs cells after 24 h treatment. Significant at *^a^p*< 0.05, *^b^p*< 0.01, and *^c^p*< 0.001 using Student’s *t*-test analysis compared to respective control. DMSO-Dimethyl sulfoxide. Data were expressed as Mean±SEM (standard error mean).

### 3-epicaryoptin induces G2/M cell cycle arrest

Considering that the most tubulin destabilizing agents could block cell division at mitosis and may lead to arrest of the cell cycle at G2/M phase. Therefore, cell cycle analysis, by using flow cytometry assay, was determined on the MCF-7 cancer cell line after 20 h treatment with the compound 3-epicaryoptin (100 and 200 µg mL^-1^). The obtained results showed that the percentage of cells in G2/M phase of the cell cycle was significantly (*p*< 0.001) increased from 12.34±0.68% for untreated control cells to 36.95±0.61 % and 24.71±0.92 % for the concentration of 100 and 200 µg mL^-1^ of 3-epicaryoptin treatment. Whereas, the percentage of cells in G0/G1 and S phase of the cell cycle decreased from 76.07±0.99 % and 9.00±0.21 % for untreated control cells to 52.92±0.37 % and 6.91±0.55 % for 100 µg mL^-1^ concentration and 67.66±1.14 % and 5.63±0.34 % for 200 µg mL^-1^ concentration respectively. Therefore, it was found that the compound 3-epicaryoptin tested against the MCF-7 cell line showed the arrest of the cell cycle at G2/M phase **(Figure 2).**

**Fig. 2.**
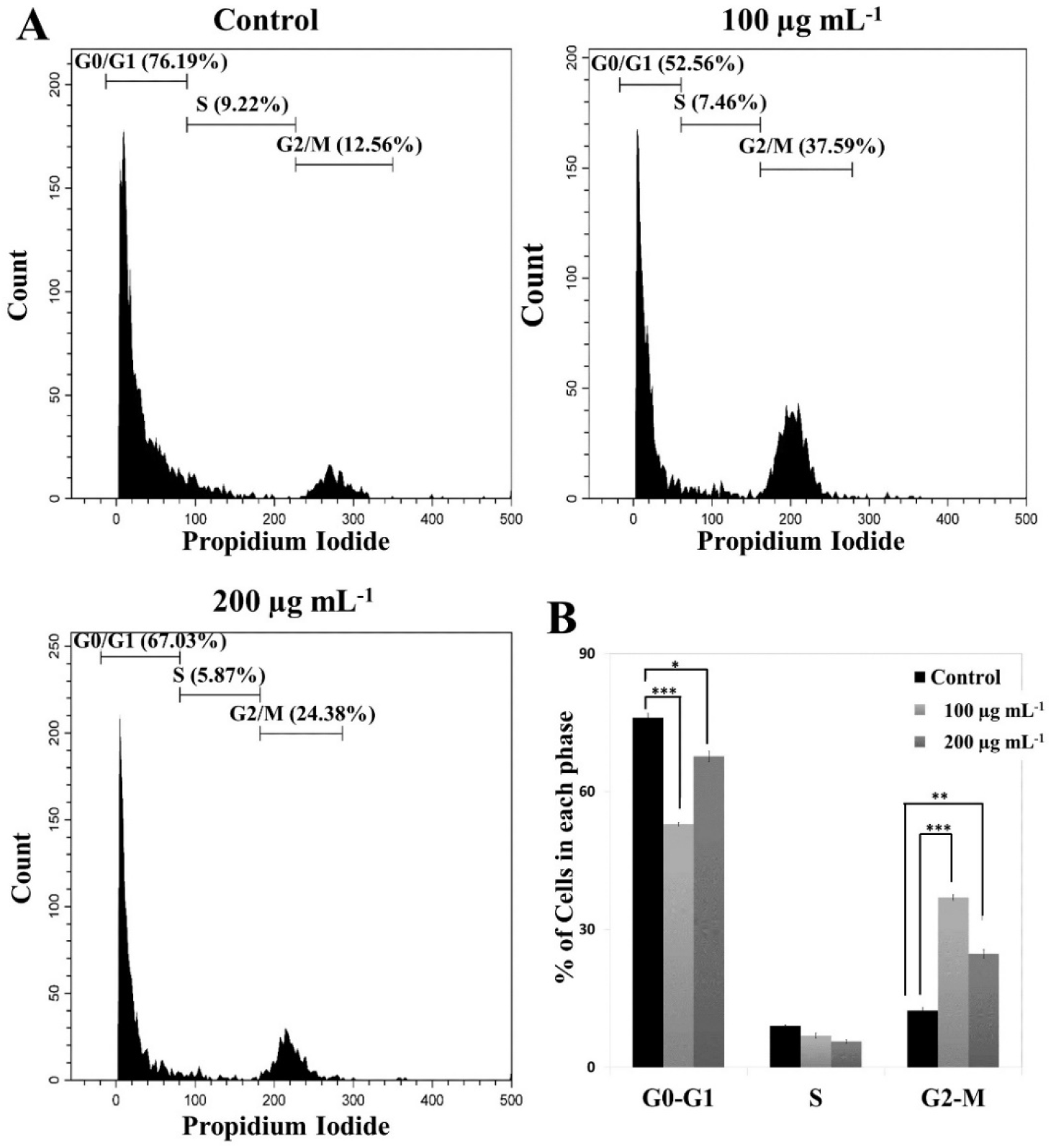
Graphical presentation of results shows the cell cycle analysis of compound 3-epicaryoptin on MCF-7 cells. (A) Represent untreated control, 100, and 200 µg mL^-1^ of 3-epicaryoptin. (B) Bar graph showing the percentage of cells in G0/G1, S, and G2/M phase. Significant at *\*p*< 0.05, \*\**p*< 0.01 and \*\*\**p*< 0.001 using Student’s *t*-test analysis compared to respective control. Data were expressed as Mean±SD (standard Deviation).

### Induction of cellular apoptosis

To examine whether the mechanism of cell death is apoptosis, we then tested the apoptosis rate in MCF-7 cells using Annexin V-FITC/PI double staining assay. The cells were treated with 100 and 200 µg mL^-1^ concentrations of compound 3-epicaryoptin or vehicle (DMSO) for 24 and 48 h and then stained with Annexin V-FITC and PI for analysis. This method divided cells into four regions corresponding to: damaged cells (Q1 region), late apoptotic cells (Q2 region), normal cells (Q3 region), and early apoptotic cells (Q4 region). Results in **Figure 3** indicated that compound 3-epicaryoptin could effectively induce apoptosis in MCF-7 cells in a time-dependent manner. The total percentage of apoptotic cells in the case of control was only 5.9% for 48 h, but the percentages were increased to 24.2% and 21.6% for 24 h, and 50.8% and 58.3% for 48 h, after treatment with compound 3-epicaryoptin at 100 and 200 µg mL^-1^, respectively. The results evidently manifested that compound 3-epicaryoptin effectively induced cell apoptosis in MCF-7 cells in comparison with the control.

**Fig. 3.**
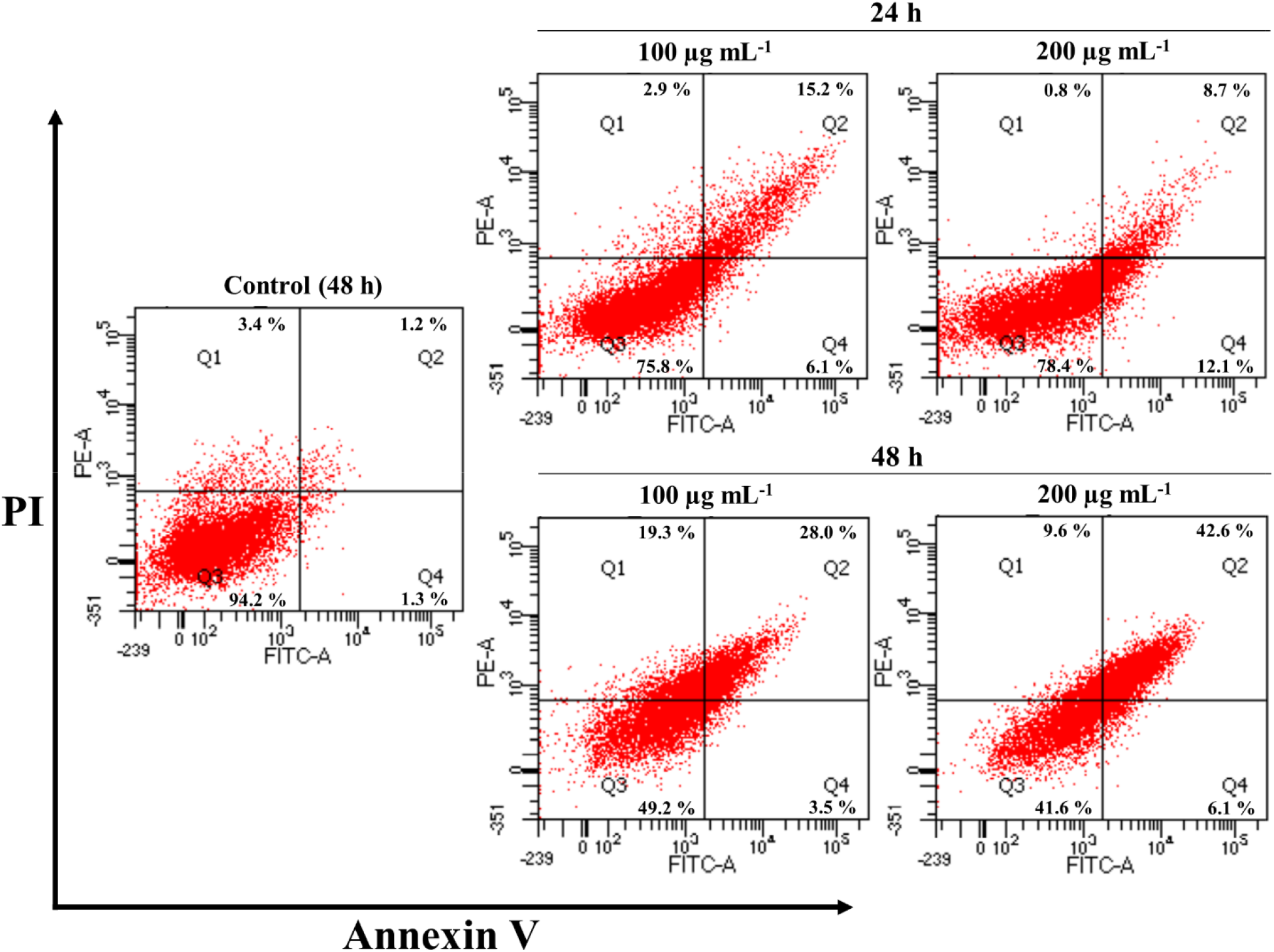
Cell apoptosis analysis of compound 3-epicaryoptin on MCF-7 cells for 24 and 48 h treatment. Cells were stained by Annexin V/PI, the induction of apoptosis was detected by flow cytometry

### Compound 3-epicaryoptin induces microtubule collapse in MCF-7 cells

For the investigation of the effect of compound 3-epicaryoptin on cellular microtubule skeleton, we performed an immunofluorescent staining assay against tubulin to determine whether 3-epicaryoptin could inhibit the microtubule dynamics in MCF-7 cells. We treated the MCF-7 cell with 100 and 200 µg mL^-1^ concentrations of 3-epicaryoptin for 20 h. As shown in **Figure 4**, the cells in the untreated control group had spindly contours and incorporated microtubule fibers. In contrast, the cells treated with 3-epicaryoptin exhibited changes in shape, and the network of microtubules became soluble and disorganized. These results indicated that the 200 µg mL^-1^ concentration of 3-epicaryoptin treated cells showed comparatively stronger depolymerizing effects than the 100 µg mL^-1^ concentration.

**Fig. 4.**
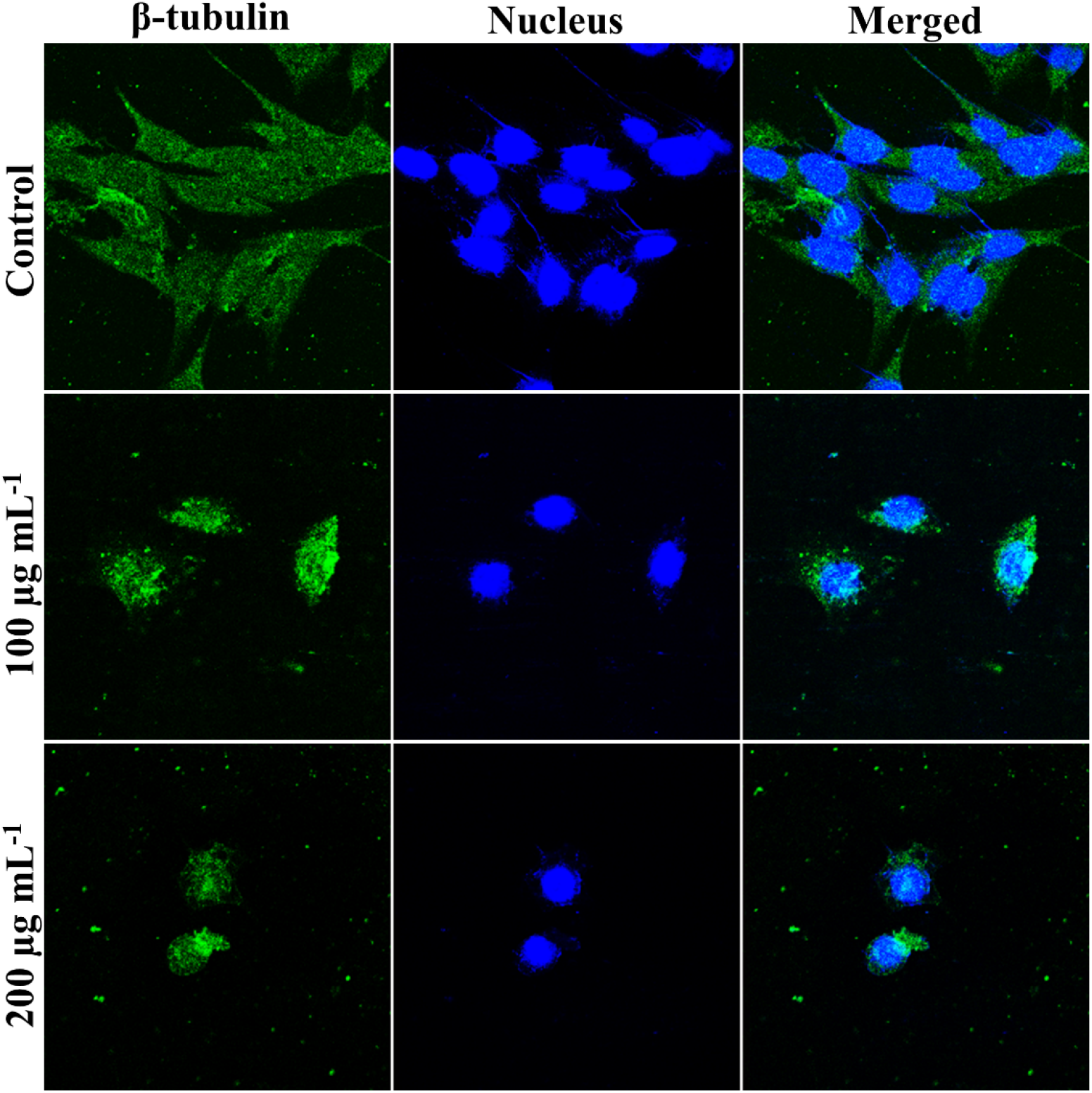
Showing 3-epicaryoptin induced microtubule depolymerizing effects in MCF-7 cells visualized by immunofluorescence and photographed using a confocal fluorescence microscope.

### Molecular docking

To examine the binding affinity of 3-epicaryoptin with tubulin, a molecular docking study was performed using Autodock Vina. The used docking protocol was validated by computing the RMSD between redocked DAMA-colchicine and its co-crystallized conformation, which was 1.076 Å in our case (**Table 1**). Generally, an RMSD value below < 2(Å) is considered acceptable [52]. As illustrated in **Figure 5**, the conformations of the original and re-docked DAMA-colchicine exhibit almost complete overlap. Which indicated that the re-docking conformation obtained through the AutoDock Vina docking protocol closely approximates the bioactive conformation of the original ligand. The analysis of docking results for 3-epicaryoptin showed that it binds to tubulin at the colchicine binding pocket (**Figure 6a and 6b**). On superimposing the docked conformations of 3-epicaryoptin and DAMA-colchicine, it was observed that 3-epicaryoptin overlapped with DAMA-colchicine (**Figure 6c**). The binding energies of DAMA-colchicine and 3-epicaryoptin with tubulin were estimated to be −8.4 kcal moL^-1^ and −9.1 kcal moL^-1^, respectively, which reveals that 3-epicaryoptin may bind to tubulin with a greater affinity than DAMA-colchicine **(Tables 1 and 2).** The LigPlot^+^ analysis indicated that the binding pocket of 3-epicaryoptin is surrounded by hydrophobic residues Ala314.B, Val313.B, Val181.A, Thr351.B, Met257.B, Asn256.B, Ala352.B, Ala248.B, Leu246.B, Leu253.B, Thr179.A, Glu183.A, Ala180.A, Lys350.B, Lys252.B, and Ser178.A. Further analysis revealed that 3-epicaryoptin showed a possible hydrogen bonding interaction with the residue Asn101 of the A chain. The distance between the positions O-29 of 3-epicaryoptin and the amide hydrogen of Asn101.A was measured to be 3.27 Å (**Table 2**; **Figure 6d**). These interactions of hydrogen bonding, together with other hydrophobic interactions, are possibly involved in stabilizing the 3-epicaryoptin in the binding pocket of tubulin.

**Fig. 5.**
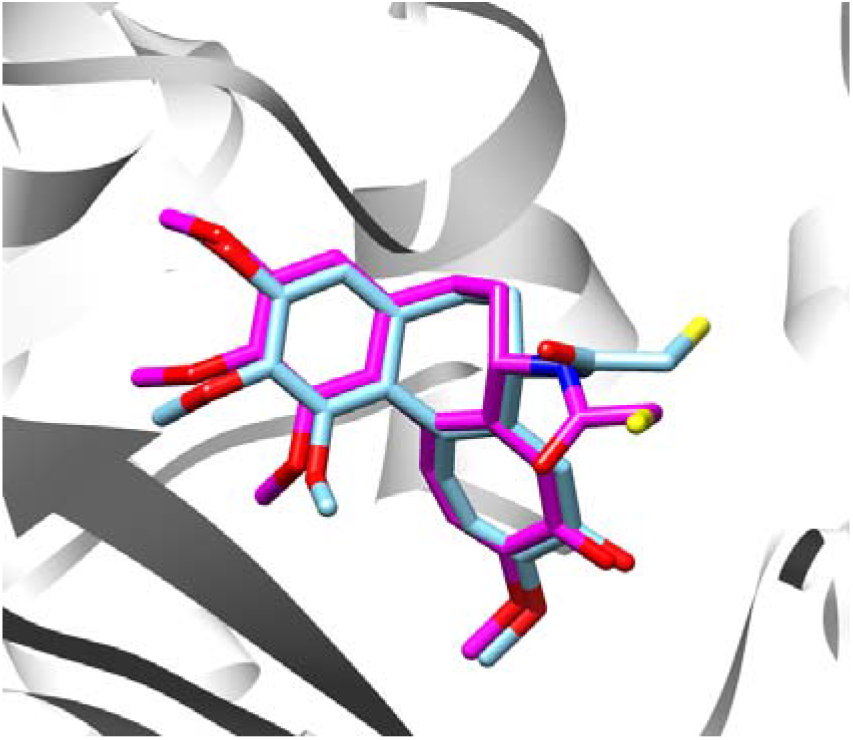
RMSD between re-docked DAMA-colchicine and its co-crystallized conformation. Coordinates of the re-docked DAMA-colchicine (sky blue) were superimposed over the X-ray crystallographically determined DAMA-colchicine coordinates (magenta) at the colchicine binding pocket.

**Fig. 6.**
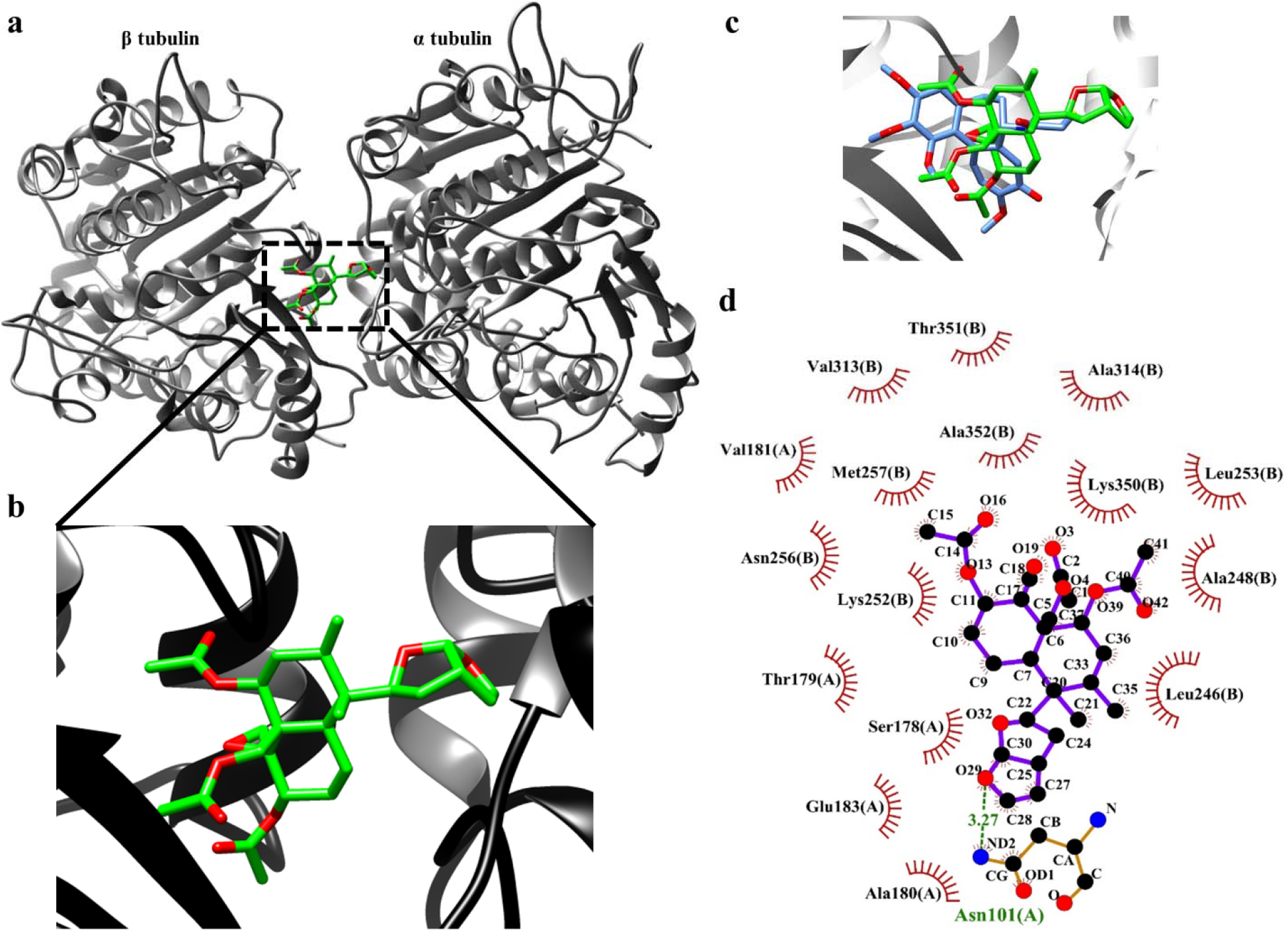
Positions of 3-epicaryoptin into the colchicine binding pocket of tubulin dimer. (a) Structural overview of αβ-tubulin with 3-epicaryoptin. (b) A close-up view of the binding pose of 3-epicaryoptin at the colchicine binding site. 3-epicaryoptin is shown in green. Red sticks represent oxygen atoms. (c) The coordinates of the docked 3-epicaryoptin (green color) were superimposed over the docked DAMA-colchicine coordinates (sky blue color) at the colchicin binding site. (d) LigPlot^+^ analysis shows the hydrogen bond (green color dot line) and hydrophobic interactions of 3-epicaryoptin with the residues of tubulin heterodimer.

**Table 1.**
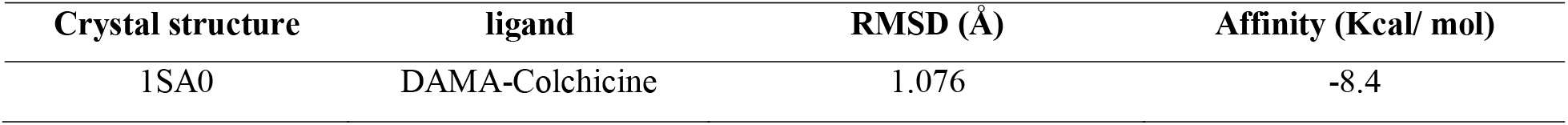
RMSD value between re-docked DAMA-colchicine and its co-crystallized conformation as well as binding affinity of re-docked conformation with tubulin.

| Crystal structure | ligand | RMSD (Å) | Affinity (Kcal/ mol) |
| --- | --- | --- | --- |
| 1SA0 | DAMA-Colchicine | 1.076 | -8.4 |

**Table 2.** Binding energy, hydrogen bonding, and hydrophobic interactions of 3-epicaryoptin with tubulin 1SA0.

| Compound | Affinity<br>(kcal mol<sup>-1</sup>) | Hydrogen bonding interactions |  | Residues in<br>Hydrophobic<br>interaction |
| --- | --- | --- | --- | --- |
|  |  | Atom interacts | Distance<br>(Å) |  |
| 3-epi | -9.1 | Asn-101-2HD....O29-3-epi | 3.27 | Ala314.B, Val313.B,<br>Val181.A, Thr351.B,<br>Met257.B, Asn256.B,<br>Ala352.B, Ala248.B,<br>Leu246.B, Leu253.B,<br>Thr179.A, Glu183.A,<br>Ala180.A, Lys350.B,<br>Lys252.B, Ser178.A |

3-epi: 3-epicaryoptin

### MD simulation analysis and prediction of binding free energy using MM-PBSA

In order to check the validity of docking results as well as to ascertain the potential binding mode, stability, and molecular interaction patterns between 3-epicaryoptin and tubulin under conditions closely resembling their natural state, MD simulations performed. The best docking pose of 3-epicaryoptin produced by AutoDock Vina was extracted and used as the initial pose for the running of a 25 ns MD simulation. The dynamic stability of the complex was assessed by calculating the RMSD values of the protein backbone and ligands along the simulation time (**Figure 7A**). Results revealed stabilization of the 3-epicaryoptin-tubulin complex structure after 10 ns, with RMSD values fluctuating between 0.35 and 0.50 nm and maintaining an average RMSD of 0.42 nm without sudden hikes or conformational changes. The ligand exhibited stable equilibrium with minimal relative fluctuation (average RMSD of 0.29 nm). Furthermore, the RMSF of Cα-atoms was calculated for the complex to identify alterations in residue flexibilities. The findings show five peaks exceeding 0.3 Å, indicating significant fluctuations in the amino acids constituting these areas. However, all these observed substantial fluctuations occur outside the αβ-tubulin interface, specifically in the loop regions, and are not influenced by the ligand (**illustrated in Figure 7B**). The structural integrity and compactness of the 3-epicaryoptin-tubulin complex were assessed by the SASA and rGy parameters. SASA results depicted in **Figure 7C** demonstrated that the complex remained compact during the trajectory, ranging between 502 and 515 nm², affirming the preservation of secondary and tertiary structures without protein unfolding. Consistent findings were observed in the rGy parameter as well. The **Figure 7D** shows no substantial variations in rGy values throughout the trajectory, confirming that the complex remained compact and the protein folding was maintained.

**Figure 7.**
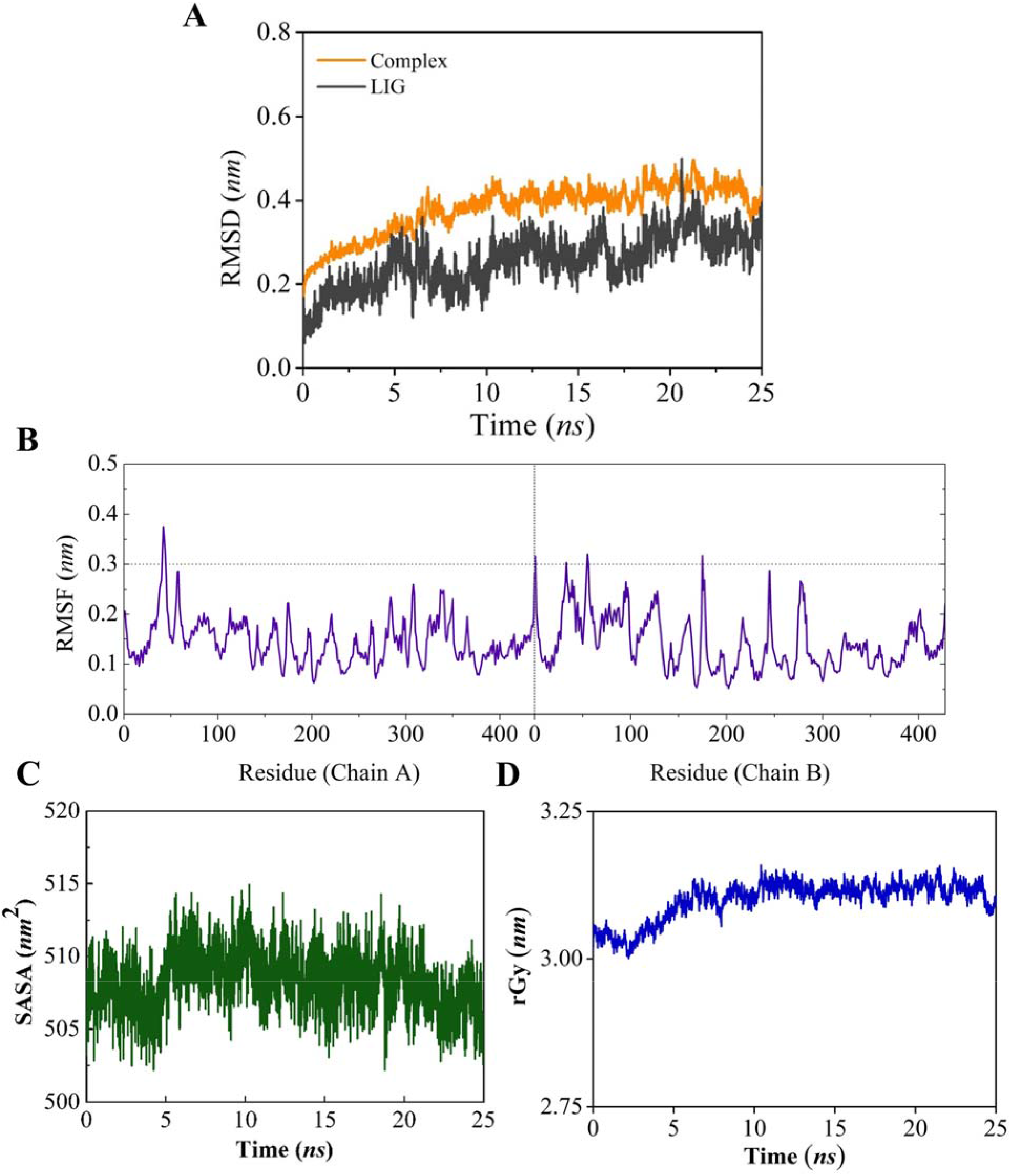
Results obtained by MD simulations of tubulin-3-epicaryoptin complex. (A) RMSD (Root Mean Square Deviation) for the complex and ligand, (B) Protein RMSF (Root Mean Square Fluctuation, (C) SASA (Solvent Accessible Surface Area) for the complex, (D) radius of gyration (rGy) for the complex.

Finally, the binding free energy was calculated using the MMPBSA method in order to study the interaction of the target residue with the ligand substructure. Different components of the binding free energy are presented in **Figure 8**. A negative value of the binding energy indicates that the ligand spontaneously binds to the protein, and the lower the binding energy, the higher the stability of the system. The estimated total binding energy of the complex was calculated as – 22.1 ± 3.4 kcal/mol, indicating good binding affinity of the ligand to the protein. The intermolecular van der Waals interaction, electrostatic interaction, and non-polar solvation terms contribute positively to the binding of the protein-ligand complex. Using the free energy decomposition, we further quantified the relative contribution of the amino acids to the compound 3-epicaryoptin (**Figure 9**). It was found that Lys252.B (−1.17 kcal mol^−1^) and Leu253.B (−2.44 kcal mol^−1^) are the amino acids that have a contribution less than −1 kcal mol^−1^.

**Figure 8.**
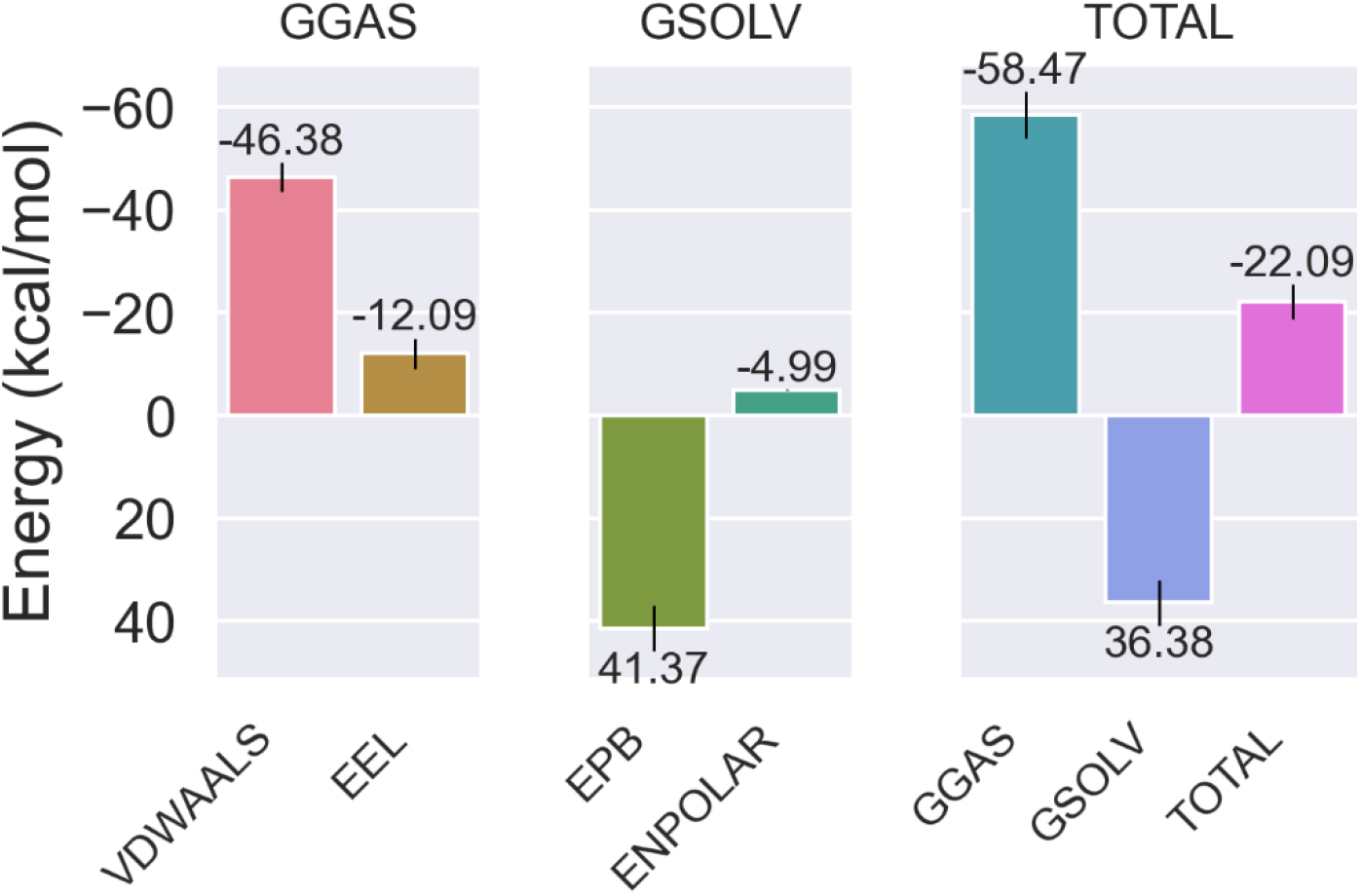
MM-PBSA energetic components and their values. Δ*E*_vdW_: van der Waals forces; Δ*E*_ele_: electrostatic energy; Δ*E*_GB_: the electrostatic contribution to the solvation free energy calculated by PB; Δ*E*_SURF_: non-polar contribution to the solvation free energy calculated by an empirical model; Δ*G*_GAS_: Gibbs free energy into a gas-phase term; Δ*G*_SOLV_: Gibbs free energy into a solvation term.

**Figure 9.**
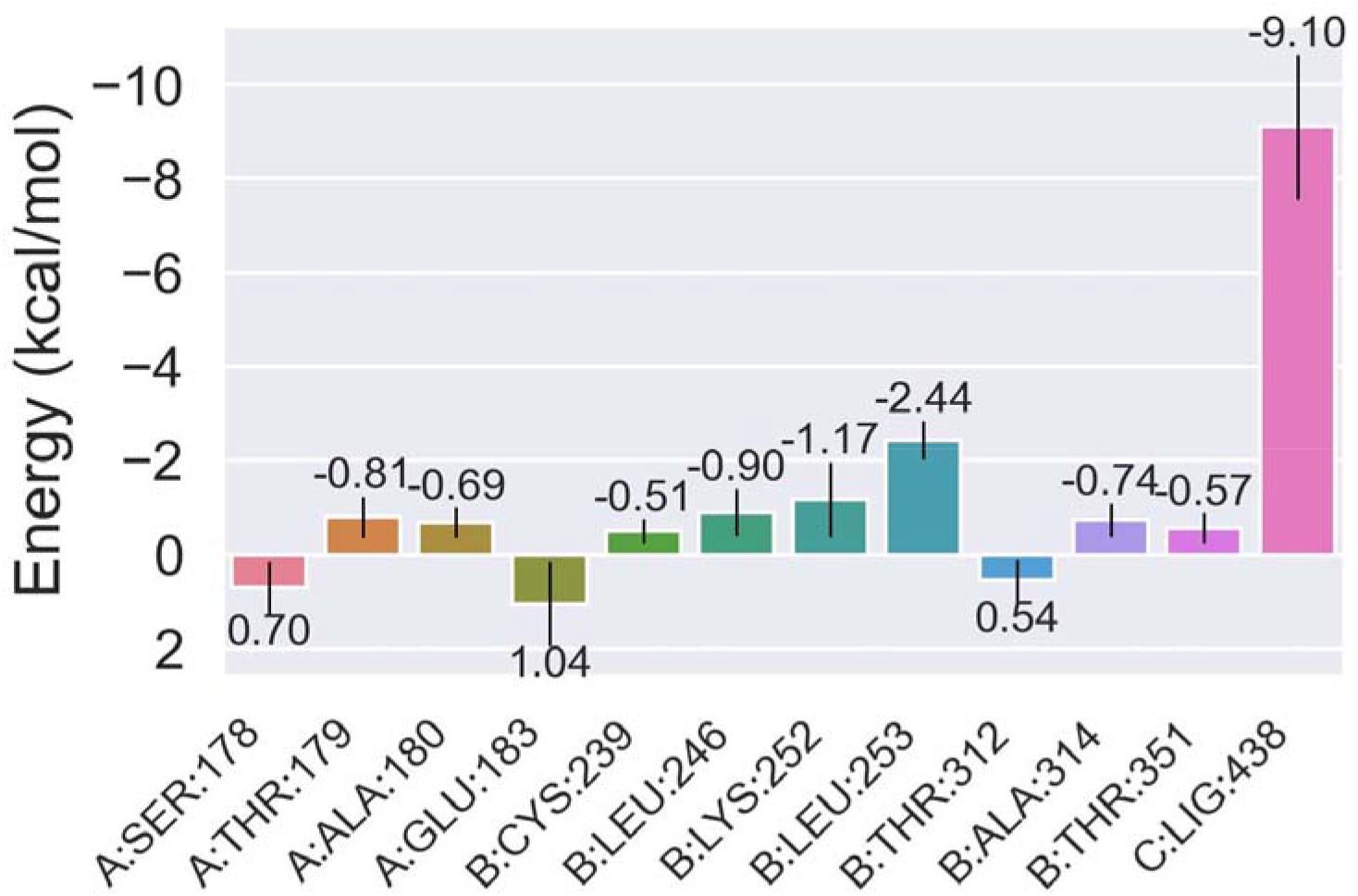
Binding free energy decomposition of the tubulin-3-epicaryoptin complex.

### ADMET profiling

The result obtained from SwissADME server is very satisfactory as 3-epicaryoptin obeyed Lipinski’s rule of five, hence possess excellent drug-likeness properties. It has high gastrointestinal absorption but does not permeate the blood−brain barrier (BBB). Additionally, it is water-soluble and possesses a favorable bioradar (**Figure 10**), making it suitable for oral administration to individuals. 3-epicaryoptin does not inhibit any cytochrome P50 enzymes and is not a P-gp substrate. A crucial log P value of 3.67, as indicated in **Table 3**, confirms its drug-like characteristics. Moreover, the total polar surface area (TPSA) of 3-epicaryoptin is also within the desirable range (**Table 3**).

**Fig 10.**
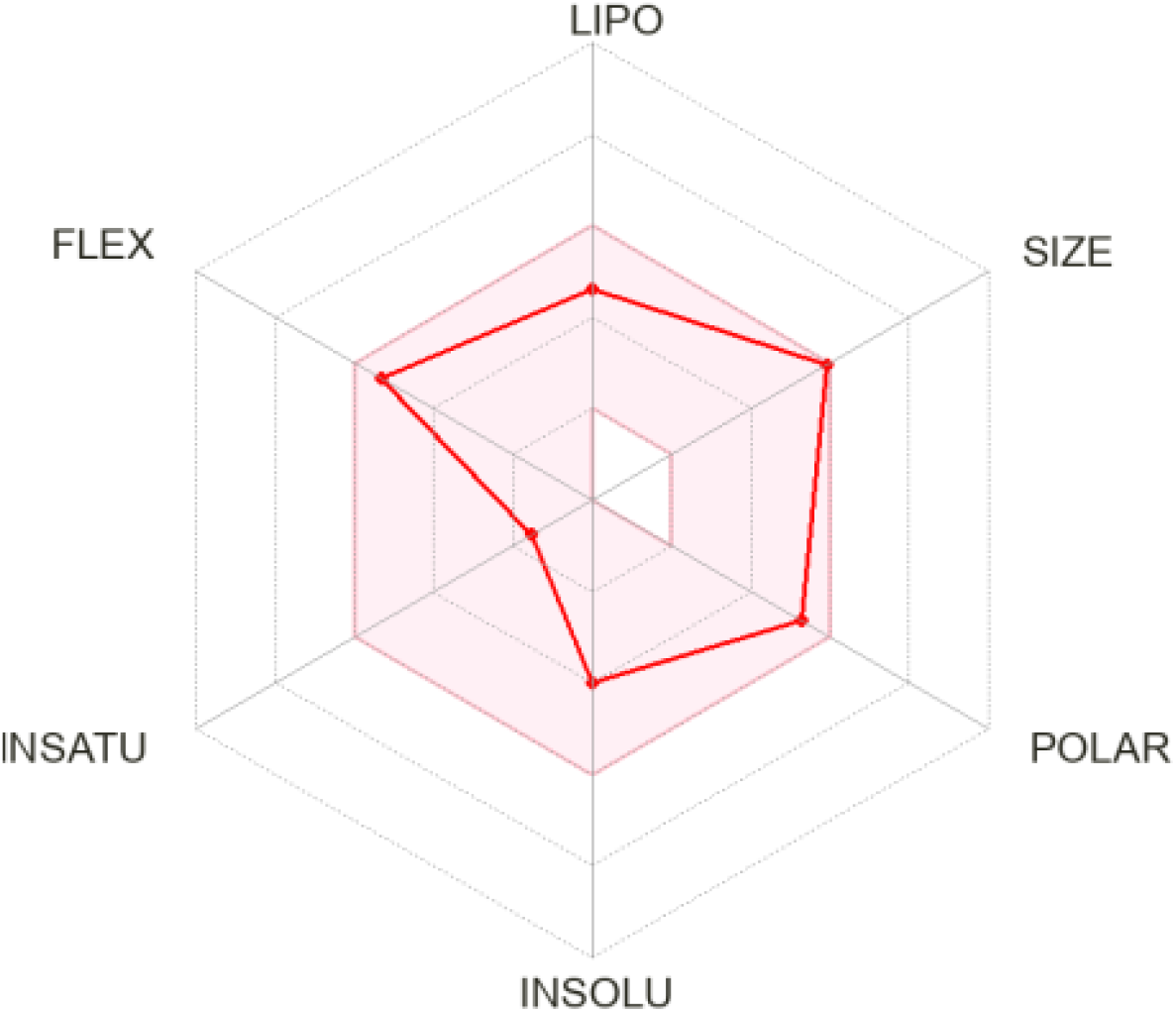
Bioradar of 3-epicaryoptin for oral bioavailability.

**Table 3.** Physicochemical, drug-Like properties, and toxicity of 3-epicaryoptin.

| physicochemical properties |  | pharmacokinetics |  |
| --- | --- | --- | --- |
| Formula | C<sub>26</sub>H<sub>36</sub>O<sub>9</sub> | GI absorption | High |
| Molecular weight | 492.56 g/mol | BBB permeant | No |
| Num. heavy atoms | 35 | P-gp substrate | No |
| Num. arom. heavy atoms | 0 | CYP1A2 inhibitor | No |
| Fraction Csp<sup>3</sup> | 0.81 | CYP2C19 inhibitor | No |
| Num. rotatable bonds | 8 | CYP2C9 inhibitor | No |
| Num. H-bond acceptors | 9 | CYP2D6 inhibitor | No |
| Num. H-bond donors | 0 | CYP3A4 inhibitor | No |
| Molar Refractivity | 122.68 | Log K<sub>p</sub> (skin permeation) | -7.49 cm/s |
| TPSA | 109.89 Å<sup>2</sup> |  |  |
| Lipophilicity |  | Drug-likeness |  |
| Log P<sub>o/w</sub> (iLOGP) | 3.67 | Lipinski | Yes; 0 violation |
|  |  | Ghose | No; 2 violations:<br>MW>480,<br>#atoms>70 |
| Log P<sub>o/w</sub> (XLOGP3) | 2.56 | Veber | Yes |
| Log P<sub>o/w</sub> (WLOGP) | 2.90 | Egan | Yes |
| Log P<sub>o/w</sub> (MLOGP) | 2.02 | Muegge | Yes |
| Log P<sub>o/w</sub> (SILICOS-IT) | 2.83 | Bioavailability Score | 0.55 |
| Consensus Log P<sub>o/w</sub> | 2.79 |  |  |
| Toxicity |  |  |  |
| Model Name | Predicted Value |  |  |
| AMES toxicity | No |  |  |
| Max. tolerated dose (human) | -0.316 |  |  |
| hERG I inhibitor | No |  |  |
| hERG II inhibitor | No |  |  |
| Oral Rat Acute Toxicity (LD<sub>50</sub>) | 4.052 |  |  |
| Oral Rat Chronic Toxicity (LOAEL) | 1.466 |  |  |
| Hepatotoxicity | No |  |  |
| Skin Sensitisation | No |  |  |
| *T. Pyriformis* toxicity | 0.286 |  |  |
| Minnow toxicity | 3.284 |  |  |

Using the pKCSM server, we conducted toxicity prediction for 3-epicaryoptin, and the outcome was very promising. 3-epicaryoptin exhibited no signs of hepatotoxicity and skin sensitization, a well as no AMES toxicity. Additionally, it displayed a Minnow toxicity score of 3.284.

## Discussion

Trends in cancer research are investigating medicines of plant origin because of their affordability and convenience, coupled with minimal adverse effects [53, 54]. In recent times, numerous compounds derived from plants have been effectively employed in cancer chemotherapy. This has sparked global interest among researchers, encouraging a thorough examination of new anticancer agents sourced from the natural environment [55–57].

Natural product 3-epicaryoptin belongs to the class clerodane diterpenoid. It’s a large class of secondary metabolites that have been studied more in recent years for their wide range of biological activities, including anticancer activity [58–62]. From previous studies, it has been well established that the compound 3-epicaryoptin has potent insect antifeedant and larvicidal activity [21, 24, 22, 23]. However, there is currently a lack of reports concerning its anticancer activity and the underlying mechanism of its inhibitory effects. Therefore, to evaluate the anticancer role of 3-epicaryoptin, we first examined the cytotoxic effect on the human BC cell line, MCF-7. As evidenced by the preliminary MTT assay, 3-epicaryoptin significantly inhibited cell proliferation at concentrations of 100, 200, and 400 µg mL-1, as depicted in **Figure 1A**. The results demonstrated a significant decrease in the viable cell percentage after 24 and 48 h of treatment, indicating the potent cytotoxic effects of 3-epicaryoptin on the MCF-7 cancer cell line [63, 57]. However, it is noteworthy to mention here that the compound 3-epicaryoptin does not harm normal human PBMCs considerably (**Figure 1B**) [64].

To investigate whether the cytotoxic effects induced by 3-epicaryoptin accompanied by alterations in cell cycle kinetics, the percentage of cells present in the different phases (G0/G1, S, and G2/M) of the cell cycle was analyzed by flow cytometric assay. The results revealed that 3-epicaryoptin at concentrations of 100 µg mL^-1^ showed the highest (36.95±0.61 %) G2/M phase cell percentage in comparison to the untreated control (12.34±0.68 % cells). These increased G2/M phase cell frequencies may specify its antimitotic effects on the MCF-7 cancer cell line [65]. However, it was observed that in the case of 200 µg mL^-1^ concentration, the G2/M phase cell frequency was again decreased (24.71±0.92 %) with an increasing (from 52.92±0.37 % to 67.66±1.14 %) G_0_-G_1_ cell frequency **(Figure 2).** This decreasing G2/M phase cell frequencies in 200 µg mL^-1^ concentration might be due to the results of an apoptotic state being increased, which was accompanied by an increase in the G_0_-G_1_ populations, and that may be determined as the major cause of 3-epicaryoptin induced inhibited proliferation in MCF-7 cells [66]. To confirm that the cells underwent apoptosis death, we performed an Annexin-V/PI staining assay of 3-epicaryoptin-treated cells. The results indicated that apoptosis was induced by 3-epicaryoptin in a time-dependent manner in MCF-7 cells, and the apoptosis rate was up to 58.3 % when treated with 200 µg mL^-1^ 3-epicaryoptin after 48 h **(Figure 3).** Hence, this present finding, consistent with previous research studies, has indicated that the G2/M phase arrest induced by 3-epicaryoptin may proceed to apoptotic cell death [67, 68].

Since, several studies was reported that the antimitotic agents exert their G2/M phase arresting effects by interfering with the cellular microtubule network [69, 70]. To determine these effects, we performed an immunofluorescence staining assay against tubulin to study whether 3-epicaryoptin could disrupt the microtubule dynamics in MCF-7 cells. As shown in **Figure 4**, 3-epicaryoptin depolymerizes the cellular microtubule network in a concentration-dependent way. Which may conclude that the 3-epicaryoptin induced microtubule depolymerizing effects are actually the mechanism underlying G2/M cell cycle arrest followed by apoptotic cell death in MCF-7 breast cancer cells [71, 72].

Numerous potent anticancer agents inhibit cell cycle progression in the G2/M phase by depolymerizing the cellular microtubule networks through direct interaction with the tubulin proteins [73, 37, 74–76]. Therefore, to further verify this hypothesis, we conducted comprehensive molecular docking and subsequent MD simulation studies to intricately illustrate the potential interactions between 3-epicaryoptin and tubulin proteins. As described in details the results section, compound 3-epicaryoptin strongly binds to tubulin between the alpha and beta tubulin interfaces, at the colchicine binding site, which involves both hydrogen bonding and hydrophobic interactions. The dynamics of the interactions, as unveiled by MD simulations, indicated the stability and compactness of the complex structure, maintaining protein folding throughout the trajectory (**Figure 7**). Further insights from the MM-PBSA method, employed to determine binding free energy, demonstrated no great variations in energy calculations, with the ligand displaying negative values for such magnitudes (**Figures 8 and 9**). These observations suggest that the systems release energy, favoring the thermodynamically favorable formation of the complex [77].

Notably, it is well-established that the anti-tubulin agents that bind at the colchicine binding pocket of tubulin hold strong antitumor or anticancer potential [47, 78, 79, 70, 80]. These agents demonstrate high anticancer efficacy and minimal off-target interactions, with many currently progressing through various phases of clinical trials for cancer chemotherapy [81, 82]. Moreover, the exploration of conjugating such agents with tumor-specific antibodies has been considered as a strategy to mitigate off-target toxicities.

So, based on the above *in vitro* experimental studies consistent with the *in silico* results, we can infer that the compound 3-epicaryoptin may act as a novel tubulin polymerization inhibitor, interfering with the tubulin at the colchicine binding site, and could display its potent antimitotic activity.

## Conclusion

The identification of intracellular molecular targets and their efficient targeting in cancer cells is the key to successful cancer therapy. Among the multiple cellular targets, the tubulin-microtubule system has been designated as one of the most classical and reliable molecular targets of the anti-cancer chemotherapeutics reported to date, especially due to its pivotal role in mitosis. Although the additional trauma caused by the systemic toxicity-related side-effects of the anticancer drugs necessitates a search for novel chemotherapeutic agents that could overcome these drawbacks. Noteworthy, this study navigates the remarkable antimitotic activity of 3-epicaryoptin against the breast cancer cell line, MCF-7. Exploration of the 3-epicaryoptin mediated G2/M phase cell cycle arresting mechanism revealed that it targeted the cellular tubulin-microtubule system and its equilibrium through binding to tubulin protein at the colchicine site, located at the interface of αβ-tubulin dimers, leading to the induction of a cell death mechanism. Hence, 3-epicaryoptin could be a promising lead compound for further development of anticancer agents through the inhibition of tubulin.

## Disclosure statement

No conflict of interest was declared.

## Acknowledgements

The authors acknowledge for the financial support of CSIR JRF-09/025(0229)/2017-EMR-I Dated: 22.08.2017, and the DST-PURSE, DST-FIST, and UGC-DRS-sponsored facilities in the Department of Zoology.

## Notes

### Competing Interest Statement

The authors have declared no competing interest.

