## Supplementary material for "3-epicaryoptin induces G2/M phase cell cycle arrest and apoptosis in human breast cancer cells by disrupting the microtubule network, an *in vitro* and *in silico* investigation"

**Table S1.** Viable cells percentage in MCF-7 cell line evaluated by MTT assay.

| **Compound** | **Concentration**  **(µg mL^-1^)** | **Viable-cells (%)** | |
| --- | --- | --- | --- |
|  |  | **24 h** | **48 h** |
| Control | - | 100 | 100 |
| DMSO | - | 98.36±1.17 | 97.65±1.73 |
| 3-epicaryoptin | 12.5 | 95.36±1.03*^a^* | 97.09±2.25 |
|  | 25 | 94.01±1.67*^a^* | 96.01±1.89 |
|  | 50 | 91.80±2.49*^a^* | 88.37±2.19*^b^* |
|  | 100 | 87.22±1.46*^c^* | 74.28±2.54*^c^* |
|  | 200 | 81.30±2.24*^b^* | 60.16±1.76*^c^* |
|  | 400 | 74.11±2.62*^c^* | 48.36±1.63*^c^* |
|  | **IC_50_** |  | **344.64** |

Significant at *^a^p*< 0.05, *^b^p*< 0.01, and *^c^p*< 0.001 using Student’s *t*-test analysis compared to respective control. DMSO- Dimethyl sulfoxide. The data represented as Mean±SEM.

**Table S2.** Viable cells percentage in PBMCs evaluated by MTT assay.

| **Compound** | **Concentration**  **(µg mL^-1^)** | **Viable-cells (%)** |
| --- | --- | --- |
|  |  | **24 h** |
| Control | - | 100 |
| DMSO | - | 99.12±0.41 |
| 3-epicaryoptin | 12.5 | 99.03±0.32 |
|  | 25 | 98.69±0.29 |
|  | 50 | 98.14±0.20 |
|  | 100 | 98.29±0.13 |
|  | 200 | 97.49±0.27 |
|  | 400 | 96.63±0.20 |

**Table S3.** Showing the percentage of cells in different (G0/G1, S, G2/M) phase by 3-epicaryoptin induced in MCF-7 cells.

| **Compound** | **Cons.**  **(µg mL^-1^)** | **Duration**  **(h)** | **Cell percentage**  **(%)** | | | | |
| --- | --- | --- | --- | --- | --- | --- | --- |
|  |  |  | **G0/G1 phase** | **S phase** | | **G2/M phase** | |
| Control | 0 | 20 | 76.07±0.99 | | 9.00±0.21 | | 12.34±0.68 |
| 3-epicaryoptin | 100 |  | 52.92±0.37*** | | 6.91±0.55 | | 36.95±0.61*** |
|  | 200 |  | 67.66±1.14* | | 5.63±0.34 | | 24.71±0.92** |

Cons.-Concentration; Significant at **p*< 0.05, ***p*< 0.01 and ****p*< 0.001 using Student’s *t*-test analysis compared to respective control. The data represented as Mean±SD.
